# Disentangling the effects of sampling scale and size on the shape of species abundance distributions

**DOI:** 10.1101/2020.01.15.900258

**Authors:** Renato A. Ferreira de Lima, Paula Alves Condé, Cristina Banks-Leite, Renata C. Campos, Malva I. Medina Hernández, Ricardo R. Rodrigues, Paulo I. Prado

**Author notes:** Naturalis Biodiversity Center, Darwinweg 2, 2333 CR Leiden, The Netherlands.

## Abstract

Many authors have tried to explain the shape of the species abundance distribution (SAD). Some of them have suggested that sampling scale is an important factor shaping SADs. These suggestions, however, did not consider the indirect and well-known effect of sample size, which increases as samples are combined to generate SADs at larger scales. Here, we separate the effects of sample size and sampling scale on the shape of the SAD for three groups of organisms (trees, beetles and birds) sampled in the Brazilian Atlantic Forest. We compared the observed SADs at different sampling scales with simulated SADs having the same richness, relative abundances but comparable sample sizes, to show that the main effect shaping SADs is sample size and not sampling scale. The effect of scale was minor and deviations between observed and simulated SADs were present only for beetles. For trees, the match between observed and simulated SADs was improved at all scales when we accounted for conspecific aggregation, which was even more important than the sampling scale effect. We build on these results to propose a conceptual framework where observed SADs are shaped by three main factors, in decreasing order of importance: sample size, conspecific aggregation and beta diversity. Therefore, studies comparing SADs across sites or scales should use sampling and/or statistical approaches capable of disentangling these three effects on the shape of SADs.

## Introduction

The species abundance distribution (SAD) is a simple yet compelling way of describing ecological communities (Pielou 1975; Chisholm and Lichstein 2009; Cheng et al. 2012). Because differences in the shape of SADs may represent changes in diversity across communities, many ecologists have tried to explain what would be the main factors shaping the SAD. There are explanations that rely on how species share niches or resources (MacArthur 1960; Tokeshi 1999) or on stochastic population dynamics (Routledge 1980; Engen and Lande 1996; Hubbell 2001). Nevertheless, it has been known for decades that the number of individuals sampled from the community (i.e. sample size) greatly affects the shape of the SADs, a phenomenon known among ecologists as the “veil line” effect (Preston 1948). In rank-abundance plots, the increase in sample size leads to less steep SADs, while in log-abundance class plots (i.e. Preston plots) this leads to more symmetrical ones (but see Nekola et al. 2008 for a discussion). Independently of the way the SAD is presented, the change in its shape is mainly due to the decline of the relative contribution of rare species in the sample (Preston 1948; Dewdney 1998; Chisholm 2007). Since the publication of Preston’s pioneer work (1948, 1962), a sampling theory was developed to deduce how the shape of SADs is affected by sample size (Bulmer 1974; Pielou 1975; Dewdney 1998). It was also shown that species aggregation and detectability patterns may also play a role in shaping SADs (Green and Plotkin 2007; Alonso et al. 2008).

More recently, the scale of sampling was suggested as an explanation for changes in the shape of SADs (Morlon et al. 2009; Šizling et al. 2009), a process that some years later was termed as scale-dependent SAD by Rosindell and Cornell (2013). Briefly, the SAD would become less steep or more symmetrical as samples from separate sites are pooled together. This trend was observed for herbs sampled over tens of meters (Wilson et al. 1998), and for trees sampled over hundreds of meters (Morlon et al. 2009). The same result was obtained for corals and reef fishes (Connolly et al. 2005) and for birds and island fishes (Morlon et al. 2009) sampled in scales of hundreds to thousands of kilometers. Indeed, computer simulations showed that the SAD do change in shape as non-overlapping samples are combined, a process that depends on the degree of species aggregation and species turnover between samples (McGill 2003; Šizling et al. 2009). Therefore, there is evidence of a sampling scale effect on the shape of the SAD, as shown for the species-area and similarity distance-decay curves (Palmer and White 1994; Nekola and White 1999).

However, previous work did not disentangle the effects of sample size and sampling scale in shaping SADs. In these studies, larger sampling scales are obtained by pooling entire samples taken from separate sites (Connolly et al. 2005; Morlon et al. 2009), leading to an increase in sampling scale *and* sample size as samples are combined (Wilson et al. 1998). Although some authors have even treated the effect of sample scale and sample size practically as equivalents (Connolly et al. 2005; Zillio and He 2010; Borda-de-Água et al. 2012), it is easy to conceive sampling designs where one can increase sampling scale without increasing sample size and vice-versa (e.g. Palmer and White 1994). One can also use randomization or rarefaction methods to tell apart the effects of sample size and sampling scale in the shape of SADs (Rosindell and Cornell 2013). Independently of the approach chosen, the control of sample size effects shaping the SAD is a crucial step in the attempt to understand the ecological processes behind community patterns (McGill and Nekola 2010; Yen et al. 2013). Surprisingly, none of the previous studies claiming an effect of sampling scale on the shape of SADs have explicitly addressed such an important issue.

Here, we disentangle the role of sampling scale and sample size in shaping SADs. Based on similar frameworks that take into account sampling intensity (Bulmer 1974; Pielou 1975; Dewdney 1998; Green and Plotkin 2007), we simulate samples of communities that preserve the properties of observed communities at a given reference scale, but that do not vary in their average sample sizes. This procedure controls for possible effects of sample size while increasing the scale of observation of the community. Following the early work of Preston (1948, 1962), our basic prediction is that sample size has a greater influence on the shape of SADs than sampling scale. Our prediction would be supported if the shape of an observed SAD matches the shape of the simulated SAD generated from higher-scale observed SADs but controlling for sample size. To assess the generality of our results, we examine our prediction using datasets of three very distinct groups of organisms (i.e. plants, invertebrates, and vertebrates) and SADs constructed using counts of individuals and biomass.

## Materials and methods

### Observed SADs

We used data from three very distinct groups of organisms sampled in the Brazilian Atlantic Forest: trees (including palms and tree-ferns), dung beetles (Coleoptera, Scarabaeidae) and understorey birds (see details in Appendix S1). Sampling was divided into four spatial scales: sample (tens of meters), habitat (one to few hundreds of meters), site (hundreds of meters to kilometers), and regional scales (hundreds of kilometers). For trees, the sample scale corresponded to 256 40×40 m permanent forest plots (total of 40.96 ha, 60 838 trees of 483 species sampled) placed in four different sites. For beetles, the sample scale corresponded to 134 pitfall traps baited with human feces (7505 beetles of 71 species captured over more than 24 000 trap-hours) placed at three sites. For birds, the sample scale corresponded to 65 sampling sites, each of them being sampled using 10 mist nets (6127 individuals and 140 species captured over a total of more than 41 000 net-hours) placed at three sites.

For all groups, habitat scale was defined by contiguous samples (plots, pitfalls and mist nets) that could be grouped based on the existence of marked environmental differences within sites, such as topographic positions, size of forest fragment, soil types or differences in disturbance history of the forest. The site scale for all groups was defined by the combination from samples taken from the same site, while the regional scale was obtained by merging all samples from all sites. Because the three groups were sampled using different methods and sampling designs, the mean number of individuals sampled at each scale, used here as a measure of sample intensity, varied among groups as well (Table S1 in Appendix S1). The population biomass of trees was obtained using allometric equations for woody species, palms, and tree-ferns. For beetles, population biomass was obtained directly from the dry weight of individuals captured in the field. For birds, the average body mass obtained from the literature was used to obtain the population biomass.

### Simulated SADs

To control the effect of sample size on SADs at different scales, we used an approach similar to parametric bootstrapping *sensu* Manly (2006). In general terms, we combined subsamples with size *n*_i_/*x* taken from *x* separate sites to create community samples at a larger sampling scales but with the same average sample size ∑*n*_i_/*x* of the lower sampling scale. In practical terms, the expected abundance of species *i* in simulated samples pooled at scale *j*, ordered from the smallest to the largest scale (*j*= 1, 2, 3, 4), was:

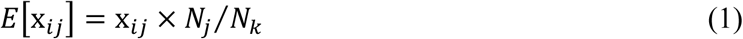

where *k* is the index for any given upper scale (*k*= 2, 3, 4); x_*ik*_ is the observed abundance of species *i* in the pooled samples at scale *k*; *N*_*j*_ and *N*_*k*_ are the observed abundances of all species at scale *j* and *k* and, thus, *N*_*j*_*/N*_*k*_ is the sampling intensity. When we combine the expected abundances of all *i* species, then the resulting SAD kept the same relative abundances of the observed SAD, but with a sample size that matches the mean size of the sample units observed at the scale below (Appendix S2). The mean size used to generate the simulated SADs varied according to the scale and group of organisms considered (Table S1). For instance, the mean sample sizes at the sample scale were 238, 58 and 94 individuals for trees, beetles and birds, respectively.

We simulated community samples by generating random draws from the Poisson distribution and the Negative Binomial (NB) distribution. Both distributions have a long tradition in ecological literature to simulate data under the assumption of spatial randomness and spatial aggregation of conspecific individuals, respectively (Pielou 1975; Brown et al. 1995; Dewdney 1998; Green and Plotkin 2007; Alonso et al. 2008). Generating random Poisson draws is similar to performing an individual-based rarefaction of the observed SAD (Connolly et al. 2005) and the comparison between the methods used here and rarefaction techniques indeed yielded similar results (not shown). On the other hand, the use of the NB adds complexity to the analysis which is related to the additional aggregation parameter (ĸ). The estimation of ĸ is generally scale and density-dependent: species with larger populations tend to have more random distributions (higher ĸ) and vice-versa (Taylor et al. 1978). In this study, we use scale-specific and species-specific values of ĸ that were estimated from the empirical data using maximum likelihood techniques. Whenever the estimation of this parameter was not possible, we assigned an arbitrary value of ĸ that was equal to the community average of this parameter at the same sampling scale. Details on the estimation of ĸ are given in Appendix S2.

Since both Poisson and NB are discrete distributions, they cannot be used directly to assess changes in species biomass distributions. Thus, the generation of simulated samples for biomass distributions was conducted in two steps. The first step was identical to the Poisson and NB draws described above, used to obtain random numbers of individuals per species. Then, for each species with *n* individuals, we sampled with replacement *n* values of biomass from the empirical biomass distribution of the same species at the same scale. Although it may not be the ideal solution, this bootstrapping procedure was very easy to implement and provided results that were appropriate enough for the purposes of this study (Appendix S2). For each group of organisms and for both measures of abundance, we draw 10000 simulated community samples.

### Observed versus simulated SADs

For each group of organisms (*i*.*e*., trees, beetles and birds) and measure of abundance (*i*.*e*., counts of individual and biomass), the observed and simulated SADs were presented using rank-abundance plot on logarithmic scales, as suggested by Newman (2005). We used relative measures of abundance in order to make the rank-abundance plots comparable among groups of organisms. Each rank-abundance plot presents only the mean of the observed and simulated SADs at each sampling scale. For instance, the observed SAD at the sample scale represents the mean SAD obtained in 256, 134 and 65 sample units for trees, beetles and birds, respectively. For the simulated SADs, plots present the mean SAD obtained from the 10000 draws for each sampling scale. For the sake of comparison, we also represented SADs using histograms of the mean proportion of species per abundances classes on a log_2_ scale (*i*.*e*., Preston plots), which are presented along with the 5% and 95% quantiles of simulated abundance values per abundance class (Appendix S3). Differently from Preston (1948), we used true octaves that are closed on the right (1, 2, 3-4, 5-8, 9-16 and so on). This is different from the binning proposal of Williamson and Gaston (2005), but the results remain essentially the same using both binning approaches (results not shown).

We used the Kolmogorov-Smirnov test to compare the observed and simulated SADs. We compared each of the 10 000 simulated SADs obtained from each reference sampling scales (site, habitat and sampling scales) with the mean observed SAD at the same sampling scale. Prior to the Kolmogorov-Smirnov test, we discretized the mean observed SAD and excluded ranks with zero individuals from both observed and simulated SADs. For biomass, we also excluded the ranks with values of relative biomass smaller than the minimum observed, which were 0.00001, 0.00005 and 0.00009 for trees, beetles and birds, respectively. Then, for each pair of observed and simulated SADs, we obtained the value of the test and the *p*-value associated with the hypothesis that the two SADs come from the same distribution. Here, we used the proportion of simulated SADs being regarded as equal to the mean observed SAD as a measure of goodness of fit. If 5% of the 10000 simulated SADs or more had a *p*-value ≥0.05, then the simulated and observed SADs were considered as being equal. The results of the Kolmogorov-Smirnov test applied using the mean observed SAD and the mean simulated SADs (i.e. only one test instead of using the mean of 10000 tests) resulted in very similar results for almost all groups of organisms and measures of abundance (Table S2 and S3). All analyses were performed in R (www.R-project.org) using the package *sads* (Prado and Miranda 2014) and *dgof* (Arnold and Emerson 2011).

## Results

All group of organisms presented very similar trends in their results using counts of individuals: the observed SADs became more concave (less linear on a log-log scale rank-abundance plots –Fig. 1) or more symmetrical (on log_2_ Preston plots – Fig. S1) as we increase the scale of observation. For SADs constructed using biomass instead of counts of individuals, the observed biomass distributions were already concave even at the smaller sampling scales (Fig. 2, Fig. S2). However, we still observed the same tendency of more concave shapes at higher scales for all groups, particularly for trees. The change of the shape of the SADs with increasing scale was more conspicuous when results are presented using Preston plots (Figs. S1 and S2) instead of rank-abundance plots (Figs.1 and 2), but the former provides a more complete representation of the response of each individual species in the community.

**Fig. 1.**
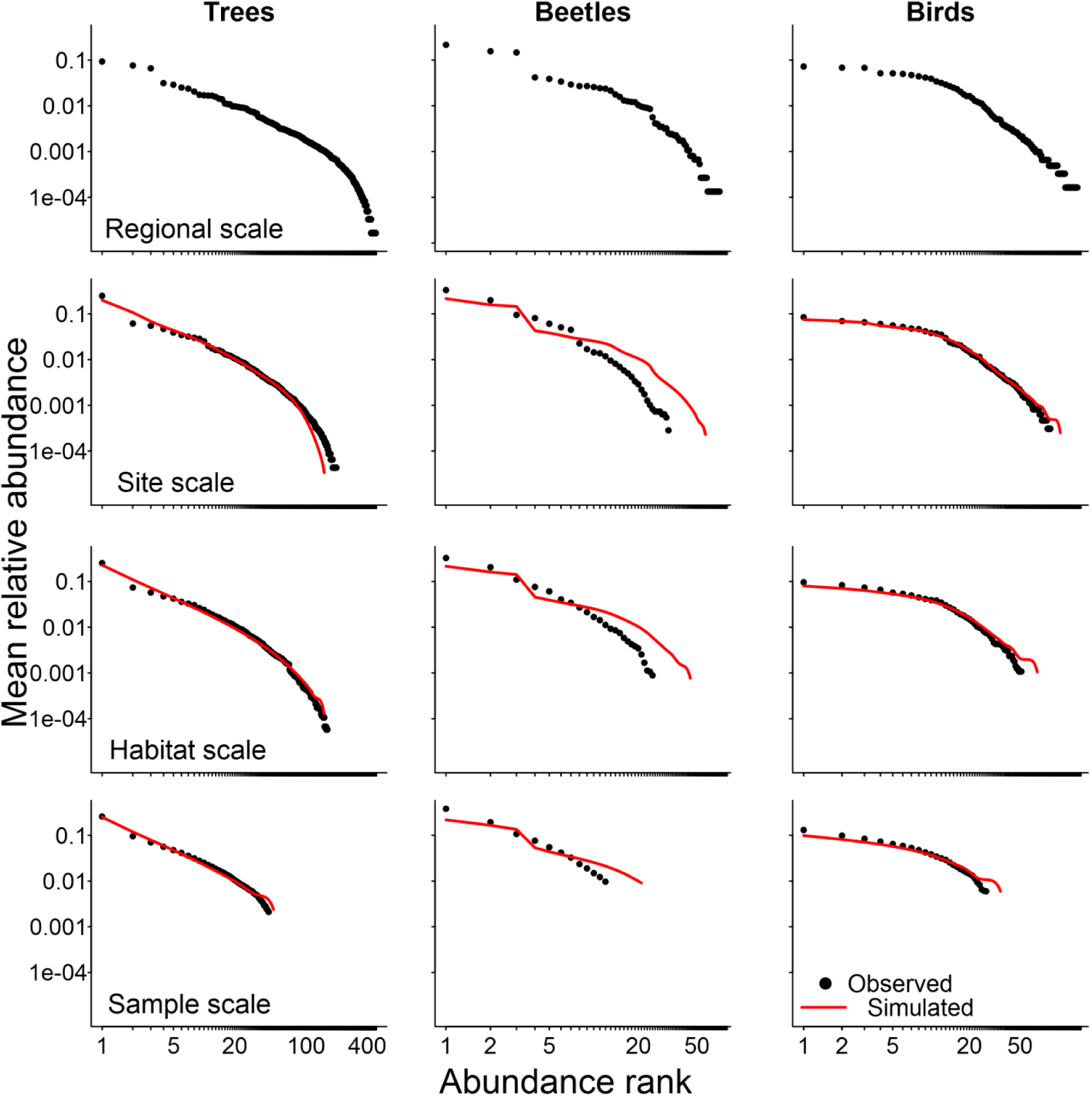
Disentangling sampling scale and sample size effects in species abundance distributions (SAD) using counts of individuals. Observed (black circles) and simulated (red lines) are the mean relative abundance of species in each rank of abundance for trees, beetles and birds at four sampling scales. For all scales except for the regional one, we present the expected values of simulated samples taken from the observed SAD at the upper scale but with the same mean sample size observed at the scale immediately below (see details on Appendix S2). Samples were simulated assuming conspecific aggregation (Negative binomial sampling) for trees and complete randomness (Poisson sampling) for birds and beetles.

**Fig. 2.**
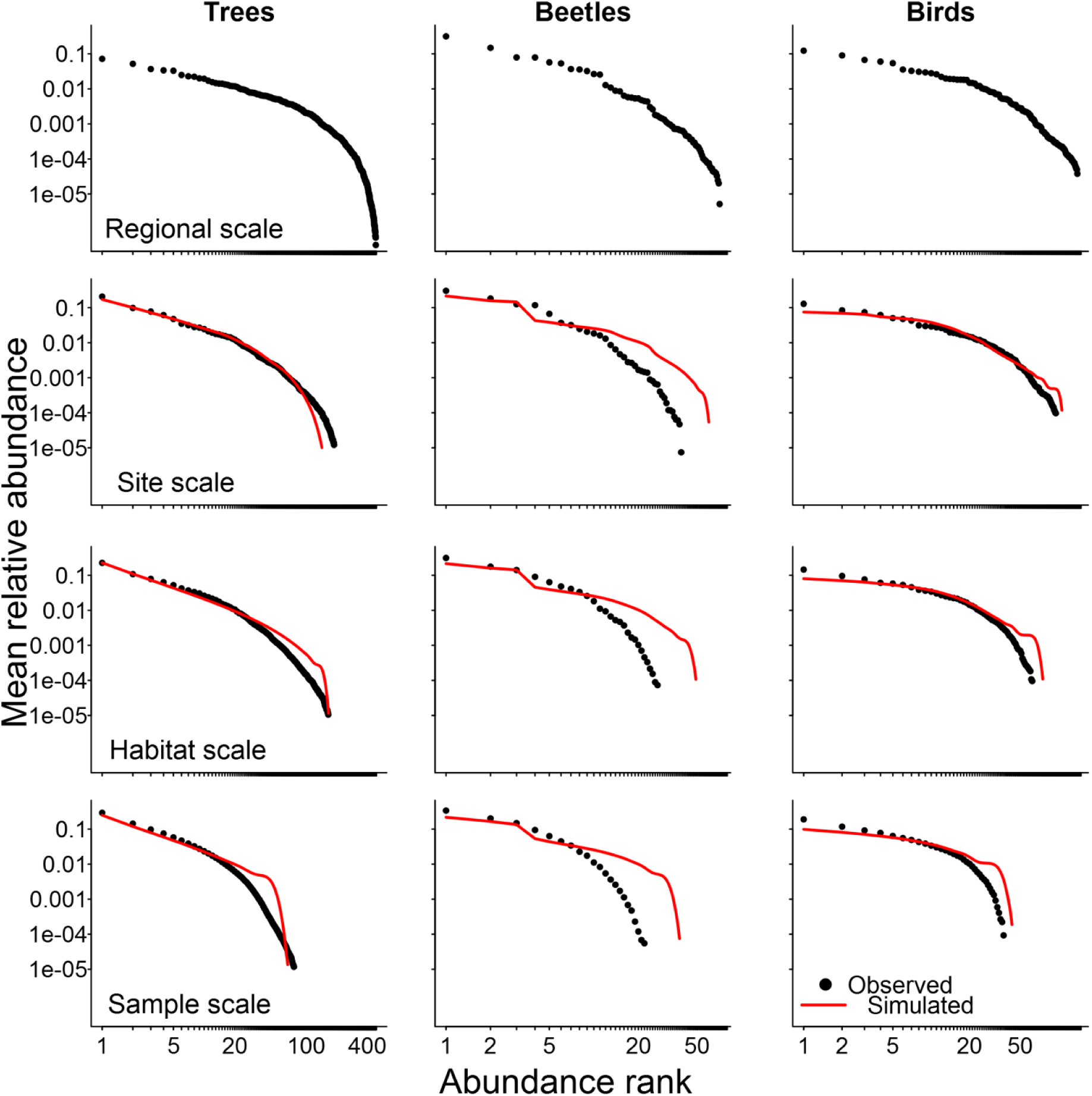
Disentangling sampling scale and sample size effects in species abundance distributions (SAD) using biomass. As in Fig. 1, observed (black circles) and simulated (red lines) are the mean relative abundance of species in each rank of abundance for trees, beetles and birds at the four sampling scales. Other details are provided in the caption of Fig. 1.

The simulated SADs presented shapes very similar to the observed ones (Figs. 1 and 2) and results were mostly the same for all groups of organisms and for both measures of abundance. The Kolmogorov-Smirnov test confirmed that simulated and observed SADs had very similar shapes in almost all possible combinations of reference sampling scales and community sample sizes (Table 1). Beetles had more deviations between simulated and observed SADs, but the Kolmogorov-Smirnov test did not suggest significant differences between their shapes. This lack of significance for this group, despite the visual differences between simulated and observed SADs (Figs. 1 and 2), may be due to the much smaller number of species of beetles in comparison to the two other groups. Simulated samples taken from Negative Binomial distribution (NB) did improve the match between the observed and simulated SADs for trees at all scales when compared to samples taken from the Poisson distribution (Fig. S3). The same was not true for beetles and birds, although the latter group did present a non-significant tendency of improvement when samples were taken from the NB. Therefore, we present simulated SADs (Figs. 1 and 2 and Figs. S1 and S2) generated by NB sampling for trees and by Poisson sampling for beetles and birds.

**Table 1.** Summary of the Kolmogorov-Smirnov tests comparing the observed and simulated SADs. Values correspond to the proportion of the 10000 simulated SADs being regarded as equal (*p*-value ≥0.05) to the mean observed SAD for each reference sampling scale. Values in bold mean that observed and simulated SADs are different (α= 5%). For each reference sampling scale (*i*.*e*., regional, site and habitat), we simulated communities using the same number of species and relative densities but with a mean sample size of the corresponding scale of simulation (*i*.*e*., site, habitat and sample).

| Simulation scale | Organism | Reference scale |  |  |  |  |  |
| --- | --- | --- | --- | --- | --- | --- | --- |
|  |  | Counts of individuals |  |  | Biomass |  |  |
|  |  | Regional | Site | Habitat | Regional | Site | Habitat |
| Site | Trees | 0.815 |  |  | 0.370 |  |  |
|  | Beetles | 1 |  |  | 1 |  |  |
|  | Birds | 0.999 |  |  | 1 |  |  |
| Habitat | Trees | 0.993 | 0.977 |  | 0.581 | 0.519 |  |
|  | Beetles | 1 | 1 |  | 1 | 1 |  |
|  | Birds | 0.995 | 0.541 |  | 0.999 | 1 |  |
| Sample | Trees | 0.982 | 0.999 | **0.000** | **0.013** | 0.754 | 0.945 |
|  | Beetles | 1 | 1 | 0.973 | 1 | 0.999 | 1 |
|  | Birds | 0.999 | 0.164 | **0.000** | 0.733 | 0.999 | 0.571 |

## Discussion

As previously found for corals, fish, trees, and birds (Connolly et al. 2005; Morlon et al. 2009), pooling samples together to increase the scale of observation does indeed change the shape of the SAD. For all groups of organisms studied here, the pooled SADs were progressively more concave down (rank-abundance plots) or more symmetrical (Preston plots). However, this change in the shape of the SAD was close to what would be expected from the increase in sample size caused by pooling sampling units. Thus, the suggested effect of sampling scale on the shape of the SAD is largely due to the well-known ‘veil line’ effect: increasing sample size decreases the probability of having rare species in the sample (Preston 1948; Dewdney 1998; Chisholm 2007). The similarity of trends found for the abundance and biomass of distinct organisms such as trees, beetles, and birds reinforces this conclusion. Besides the indirect effect of sample size, the shape of SADs is also influenced by conspecific aggregation, which tends to increase the number of both rare and very common species in the sampled SAD (Green and Plotkin 2007). In our study, aggregation was important for trees (Fig. S3), confirming that tropical tree species are often spatially aggregated (Condit et al. 2000; Capretz et al. 2012). This also suggests that the role of conspecific aggregation in shaping SADs may be more important for sessile organisms, at least for the spatial scales considered here. And if we consider that the patterns of species aggregation depend on the scale considered (Taylor et al. 1978; Fortin and Dale 2005), then conspecific aggregation may even be regarded as an indirect effect of the sampling scale.

Although sample size is the preponderant effect on the SAD, we found some deviations between observed and simulated SADs that could not be attributed to sample size or conspecific aggregation. Thus, there is evidence of a sampling scale effect, which is related to the beta diversity of the metacommunity (Šizling et al. 2009). Beta diversity depends on the correlation of species abundances across communities, which in turn defines the joint SAD of two or more communities as a multivariate probability distribution (Engen et al. 2002; Sæther et al. 2013). Taking two communities as a simple example, there is a small probability that a given species is very abundant or very rare in both samples when beta diversity is high (negative correlation, inset plot in Fig. 3a). This probability is higher when beta diversity is low (positive correlation, inset plot in Fig. 3b). Therefore, when we combine samples the resulting SAD will have larger variance as beta diversity increases, because the proportion of rare species in the pooled SAD increases. In our study, beta diversity was higher for beetles at all sampling scales, which may explain the deviations between observed and simulated SADs found for this group. Moreover, conspecific aggregation also increases the variance in the correlation between samples and of the pooled SAD (Figs. 3c and 3d – see Fig. S4 for the same results presented using Preston plots). Unlike the effect of sample size and species aggregation, which can be efficiently handled using sampling theory, beta diversity is related to ecological and evolutionary processes, such as habitat partitioning, dispersal limitation, biogeography and disturbance history. Since these processes are hardly perceived at very small scales (e.g. Cheng et al. 2012), beta diversity will probably affect the shape of SADs at larger sampling scales.

**Fig. 3.**
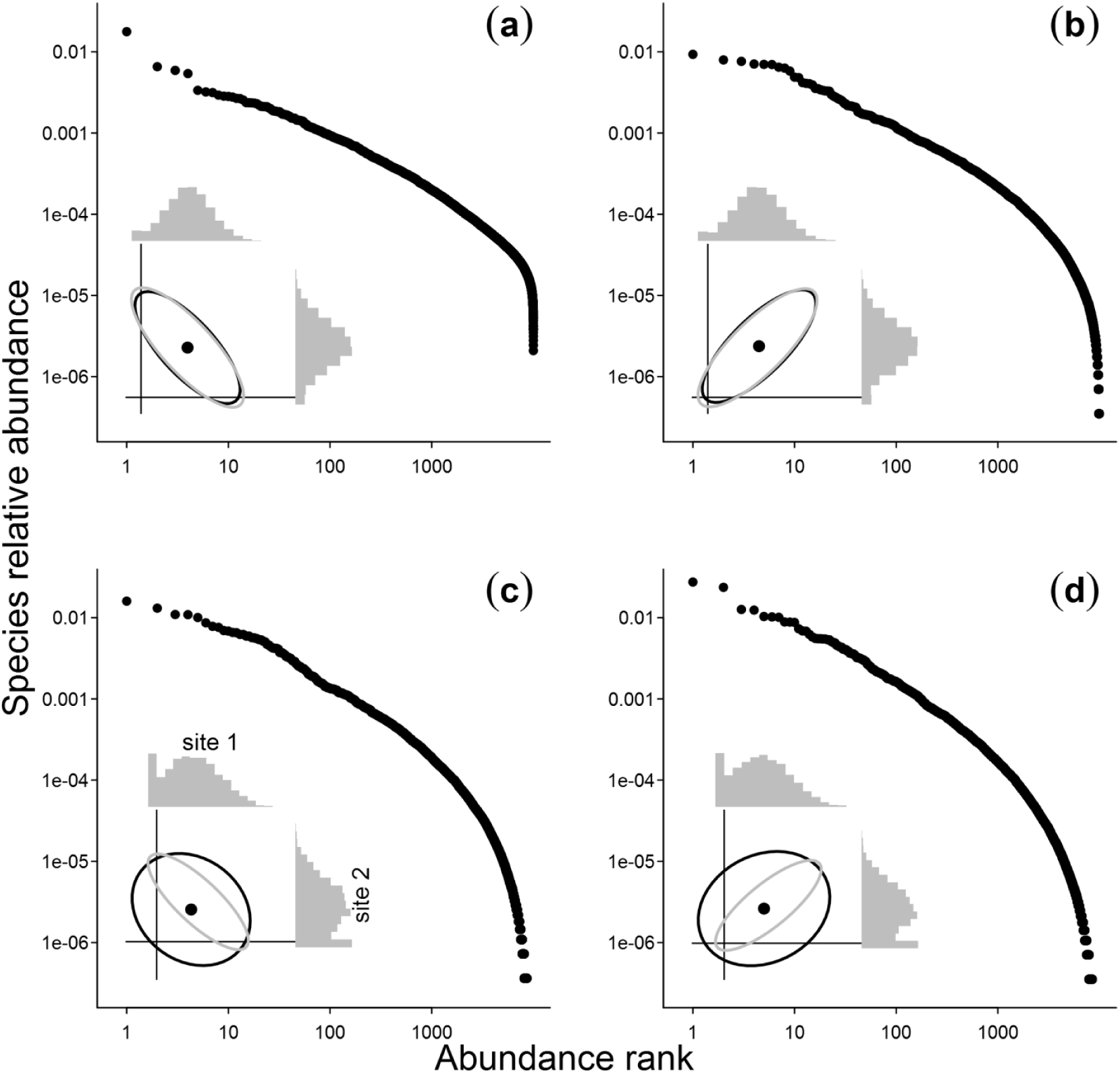
The influence of conspecific aggregation and of the correlation of species abundances among communities (beta diversity) in the shape of the SAD, while combining samples to increase the sampling scale. Here, we exemplify this influence by pooling together two samples taken from a bivariate Lognormal distribution with the exact same parameters and controlling for possible sample size effects. We present the resulting rank-abundance plots (on log-log scale) from the combination of sites where individuals of all species are randomly distributed and beta diversity is high (a) or low (b) and when there is conspecific aggregation with high (c) or low beta diversity (d). In each panel, the inset plot shows the ellipses that encompass 95% of the bivariate distributions and the marginal Preston plots of the sample SADs. Gray ellipses delimit the joint distribution of abundances in the communities and black ellipses delimit the distribution of abundances in the samples.

Engen et al. (2002) advanced the use of multivariate distributions to describe joint SADs. By assuming no conspecific aggregation and a Lognormal underlying SAD, they used a bivariate Poisson-Lognormal distribution to describe joint SADs where (i) communities can have different underlying SAD parameters, (ii) with different correlation between species abundances, (iii) which will present a sampling size effect on the SAD of the pooled samples. Sampling decreases the correlation between species abundances, an effect that is stronger if there is conspecific aggregation (gray and black ellipses in the inset plots of Fig. 3 and Fig. S4). Parametric multivariate distributions are still hard to fit, especially if we want to take into account spatial aggregation. So far, the existing investigations are based on simulations and they have shown that the similarity between sampled communities is important (McGill 2003; Šizling et al. 2009). However, these investigations were not fully parametric and/or they were based on presence-absence similarity indexes. Here we show that the simulation from parametric multivariate distributions with an explicit correlation between species abundances is feasible, as exemplified in Fig. 3 (see Appendix S4 for details), and thus one could use simulation-based methods of inference (Hartig et al. 2011) to fit and compare multivariate SADs across space and time. We also present different ways of presenting pooled SADs from samples taken at different scales (Appendix S5).

As found by Connolly et al. (2005) and Morlon et al. (2009), SADs built using biomass instead of counts of individuals are more symmetrical in Preston plots, a phenomenon termed ‘differential veiling’ by Williamson (2010). Here, we attribute this difference to the fact that biomass SADs are essentially the distribution of sums of biomass values (i.e., rare species will not necessarily have the lowest biomass ranks). If we take biomass as random variables, then the higher symmetry of the biomass SAD can be the product of a weak variant of the Central Limit Theorem: the sum of independent variables will tend to a normal distribution (Johnson et al. 1994). It is reasonable to assume that biomass values are independent within species, but biomass distributions are probably not identically among species. Thus, biomass SAD will tend to a more symmetrical distribution, but more slowly and deviating from normality. This is a different explanation than those provided by Williamson (2010) and Anderson et al. (2012), both relying on differential consequences of sampling individuals (discrete) and biomasses (continuous variable). Ecologically speaking, the higher symmetry of the biomass SAD suggests that the dominance in ecological communities is lower than previously thought. If biomass expresses better the partitioning of resources among species (Saint-Germain et al. 2007), a few but large individuals could suffice to maintain populations, especially regarding size-structured organisms. And since growth is more frequent than births and deaths, the biomass SADs is more variable and may be more effective to capture changes on community structure (Morlon et al. 2009; Anderson et al. 2012).

We conclude that sample size has a much greater influence than sampling scale on the shape of SADs. Although sample size was the leading effect, we propose that the shape SADs is influenced by three factors, in decreasing order of importance: sample size (Preston 1948, 1962), conspecific spatial aggregation (Green and Plotkin 2007) and beta diversity (McGill 2003; Šizling et al. 2009). The changes related to the first two factors can be taken into account using an approach based on univariate distributions, while the third may require the use of multivariate distributions (Engen et al. 2002; Sæther et al. 2013). Analytical solutions for such distributions may be hard to find or to deal with, and simulation-based methods of inference offer a more feasible alternative for most ecologists. The present study and some previous work (McGill 2003; Šizling et al. 2009) showed how computer simulations can address the ‘process and observation layers’ (Royle and Dorazio 2008) to disentangle the effects of sampling scale and sample size on the shape of SADs. Future studies comparing results with different underlying SADs, species aggregation patterns and beta diversity scenarios are welcome to refine our understanding of the effects of such factors on SADs and how such effects interact with existent hypotheses that explain the shape of SADs based on species dispersal distances (Chisholm and Lichstein 2009), species-range distributions (Brown et al. 1995; McGill and Collins 2003) or species strategies (e.g. core-occasional species hypothesis – Magurran and Henderson 2003). In addition, there is still uncertainty if these hypotheses apply similarly to numerical abundance and biomass (Morlon et al. 2009; Henderson and Magurran 2010). Thus, once the effects of sample size and species aggregation have been taken into account, there is still plenty of room to study the role of ecological determinants of SADs (McGill and Nekola 2010; Yen et al. 2013).

## Supporting information

Supporting R Codes

## Acknowledgments

We thank Vinícius C. Souza and Fernando Z. Vaz-de-Mello for the identification of tree and beetle species, respectively. We also thank Hélène Morlon and Jerôme Chave for their valuable suggestions. Data collection was supported by the grant 1999/09635-0 and 2013/50718-5, São Paulo Research Foundation (FAPESP) and by CNPq/BMBF (process 690144/01-6). RAFL was supported by CAPES and by grant 2013/08722-5, São Paulo Research Foundation (FAPESP). PIP was supported by CNPq (process 303878/2008-8) and by grant 09/53413-5, São Paulo Research Foundation (FAPESP).

## Author contributions

RAFL, PAC and PIP conceived and designed the study. PAC, CB-L, RCC., MIMH and RRR gathered and validated the data. RAFL and PIP analysed the data and wrote the draft of the manuscript. All authors participated in the review and editing of the final manuscript.

