## Supporting Information for "Disentangling the effects of sampling scale and size on the shape of species abundance distributions"

**This PDF file includes:**

**Appendix S1-S5**
**Figures S1-S5**
**Tables S1-S3**

**Appendix S1**. Details on the sampling methods adopted for the three groups of organisms.

Trees were surveyed using permanent forest plots placed in four forest types in São Paulo State, namely the savanna, seasonal, white-sand, and rain forests. The two first sites are inland and 75 km apart from each other. They are ca. 300 km apart from the other two coastal plots that are 100 km apart from each other. Each plot has 320×320 m (10.24 ha) and was subdivided into 64 subplots of 40×40 m, regarded here as sample units (total of 256 subplots). This subplot size was chosen to avoid samples with low number of individuals and to avoid spatial correlation of community properties between samples, which ranged up to 20-30 m (R.A.F. Lima, unpublished data). All trees, palms, and tree ferns with stem girth at breast height (1.3 m) ≥ 15 cm were tagged, measured and identified to species. Individual biomass was obtained using allometric equations for moist forests based on tree diameter and species-specific wood density (Chave *et al.* 2005). For *Euterpe edulis*, a very abundant palm, and tree ferns (Cyatheaceae) we used specific formulas given by Wendling (1998) and Tiepolo *et al.* (2002), respectively. The results refer to the second censuses conducted in 2004/2005 (total of 60 838 trees and 483 species).

Dung beetles (Scarabaeidae) were surveyed at different three sites, namely Serra do Japi (São Paulo State), Lagoa do Peri and Campos Novos (both in Santa Catarina State). At Serra do Japi sampling was conducted between 1997 and 1998 in six locations, using four pitfall traps per location (Hernández & Vaz–de–Mello 2009). At Lagoa do Peri sampling was performed between 2007 and 2008 in two different locations, using five pitfalls per location (Condé 2008). At Campos Novos, sampling was carried out in 2011 in 20 different forest fragments, using five pitfall traps per fragment (Campos & Hernández 2013). At the first two sites traps were placed along transects at intervals of 50 m, while at Campos Novos we used intervals of 10 m. Dung beetle collections were undertaken with traps baited with human feces and left open for two days every month. Thus, individuals captured monthly during one year (576 trap-hours) were assembled to compose the sampled beetle community of each trap. The exception was Campos Novos, where sampling was conducted only once (48 trap-hours). Each trap was regarded as separate sample unit (total of 134 traps). All beetles were identified to species, oven-dried at 40-45ºC for at least 48 hours and weighted to obtain their dry weight, used here as a measure of biomass. In the three sites, sampling resulted in a total of 7505 beetles and 71 species captured over more than 24 000 trap-hours.

Understorey birds were surveyed between 2001 and 2007 at three sites in São Paulo state, Brazil. Sites were 50-150 km apart from each other and varied in their percentage of forest cover (10, 30 and 50%), each of them being composed by forest fragments of different sizes (2 to 150 ha) and by more continuous forest areas (Banks-Leite *et al.* 2012). Sampling was conducted using mist nets (12 m long, 2.5 m high, 18 mm mesh) positioned inside different forest fragments and at different areas of continuous forest areas, comprising a total of 65 samples in the three sites (53 in forest fragments and 12 sites in continuous forest areas). In each of these sites, we placed 10 mist nets that remained opened twice in the dry season and twice in the wet season (average ± standard deviation of 637 ± 77 net-hours per site). Therefore, birds captured over one year were assembled to compose the sampled bird community of each net. All birds captured were tagged with an aluminum band, identified to species and released in the vicinity. The mean body mass in grams was obtained from the literature to obtain the total biomass of each species. A total of 6127 individuals and 140 bird species were captured in the three sites over more than 41 000 net-hours.

As stated in the main text of this study, the three groups of organisms were sampled in four different spatial scales: sample, habitat (within-site), site, and regional scales. The mean number of individuals sampled at each scale, which can be used as a measure of sample intensity, varied among groups of organisms and was higher for tress followed by beetles and birds (Table S1). The division of habitats for beetles and birds was very straightforward since sample units were intentionally placed on separate types of environments such as valley bottoms, slope or ridge tops, and forest fragments of different sizes. Forest fragments were divided into very small (< 2 ha), small (2 to 9 ha), medium (12 to 40 ha), large (45 to 150 ha) and very large (> 1000 ha). For trees, which were sampled using contiguous 40×40 m plots, habitat scale was defined first by classifying each of the four 20×20 m quadrats that comprise the 40×40 m plots into one habitat class, such as soil types or disturbance histories. Then, we classified each of the 40×40 m plots by counting the number of quadrats in each habitat class. If the 40×40 m plot had more than three quadrats in one habitat class, the plot was classified accordingly. If the plot had two quadrats of different classes, then the plot was arbitrarily assigned to one of the two classes of habitat, giving preference to the habitat class with smaller total area inside the site. This procedure was adopted to avoid an excessive number of habitat classes and although arbitrary it was necessary for the classification of only 19 of the 256 40×40 m plots. In the rainforest plot, soil patches with small sizes (covering two or less 40×40 m plots) were not considered as separate habitats and were included in existing habitat classes that were more similar in terms of environmental properties.

**Table S1.** Definition of the regional, site, habitat and sample scales used in this study and the respective number of sampled communities at each scale. The mean number of individuals sampled at each scale (Mean[$n_i$]) is given for each scale.

| **Group** | **Sampling scale** | | | |
|---|---|---|---|---|
| | **Regional** | **Site** | **Habitat** | **Sample** |
| Trees | São Paulo | Savanna | Disturbance history (×4) | Subplots (×64) |
| | | Seasonal forest | Soil type (×4) | Subplots (×64) |
| | | White-sand | Soil type (×3) | Subplots (×64) |
| | | Rain forest | Soil type (×4) | Subplots (×64) |
| Mean[$n_i$] | – | 15210 | 4055 | 238 |
| Dung beetles | São Paulo and Santa Catarina | Serra do Japi | Topographic position (×6) | Pitfall traps (×24) |
| | | Lagoa do Peri | Topographic position (×2) | Pitfall traps (×10) |
| | | Campos Novos | Size of fragment (×3) | Pitfall traps (×100) |
| Mean[$n_i$] | – | 2502 | 682 | 58 |
| Birds | São Paulo | Ribeirao Grande | Size of fragment (×4) | Mist nets (×21) |
| | | Caucaia do alto | Size of fragment (×4) | Mist nets (×21) |
| | | Tapiraí | Size of fragment (×4) | Mist nets (×23) |
| Mean[$n_i$] | – | 2042 | 511 | 94 |

**Appendix S2**. A basic tutorial to generate the random draws based using the observed community properties from the Poisson and the Negative Binomial distributions. Codes are given in language R (http://www.r-project.org).

Under the assumption of spatial randomness of species, generating draws to simulate community samples with decreasing sample sizes is quite straightforward, especially when dealing with counts of individuals. Suppose that we have a given community based on which we want to generate simulated samples and let this community be represented by a vector of species abundances *V*. Because each element of *V* is simply the abundance of each species present in this community, the length of *V* is the community richness while the sum of its elements is the community size. As cited in the text, the Poisson distribution is often used to mimic a sampling process where there is an assumption of spatial randomness. Therefore, the generation of random draws with sample intensity *a* based on an arbitrary community can be performed using the following R codes:

```
# Creating an arbitrary vector of abundances V, with 20 species and 300
#individuals:
      V <- c(76,5,30,3,1,16,9,1,2,42,13,14,1,1,8,53,3,18,3,1)
      S <- length(V)
      N <- sum(V)
# Setting sample size to one third of V, i.e., 100 individuals:
      a <- 100/N
# Poisson draws with the same community properties but smaller size:
      v <- rpois (n= S, lambda=V*a)
```

Note that *v* has on average the same relative abundances of *V* but the sum of the vector *v* is on average *a* times smaller than *N*. In addition, *v* has species with abundance zero, meaning that there are species present in the original community that were not ‘sampled’. Thus, the species richness of *v* will be smaller than *V* (about 14.5 ± 1.5 species in the example above). If we replicate this procedure many times, we can establish what would be the mean shape of the species abundance distribution (SAD) if the original community had the same community properties (*i.e.* species richness and relative abundances) but a sample size *a* times smaller.

If species are aggregated, we can use the Negative Binomial (NB) distribution to assess changes in the shape of the SAD as a function of sample size. Differently than the Poisson distribution, which has only one parameter, the NB has an additional overdispersion parameter κ that measures the amount of species aggregation (Bolker 2008). Thus, to generate random draws from the BN distribution we use the same vector of species abundances *V* but we have to provide values of species aggregation as well:

```
# Creating arbitrary values of k for each species
      k <- c(0.5,0.15,1,0.1,0.1,0.15,0.1,0.1,0.2,1.5,0.3,0.1,0.05,
      0.05,0.8,0.5,0.1,0.9,0.1,0.05)
# NB draws with the same community properties but smaller size:
      v <- rnbinom (n= S, mu=V*a, size=k)
```

Once again, note that sum of the vector *v* is on average *a* times smaller than *N* and that it also has species with abundance zero. But since aggregation decreases the expected number of species in samples (Hughes 1986, Colwell *et al.* 2004), the mean richness of *v* generated using the NB distribution (8.5 ± 1.7 species) is smaller if compared to the Poisson distribution.

In the analytical framework used in the manuscript, the vector *V* was defined based on the observed SAD of each reference scale (regional, site, and habitat scales) and the sample intensity *a* was defined based on the mean sample size *n* observed at each target scale (site, habitat, and sample scales). For instance, when the regional scale was used as a reference *V* was defined as a vector containing species from all sites and their overall abundances, and the values of *a* for the target scales were defined as $a_{site}= n_{site}/N$, $a_{habitat}= n_{habitat}/N$, and $a_{sample}= n_{sample}/N$. Since the higher scale of observation in our study is the regional one, than values of *N* in the manuscript represent the total of individuals sampled at this scale (60 838, 7505 and 6127 for trees, beetles and birds, respectively).

Although the simulated samples from the observed SAD at regional scale could be generated right away, the simulated samples from the observed SADs at site and habitat scales had to be generated in two levels. First, we simulated separate samples based on the observed community properties of each individual site or habitat. These samples were generated preserving the observed relative abundances of each site or habitat, but using a common sample size for all habitats or sites. Again, sample sizes were defined based on the mean sample sizes observed at each scale and for each group of organisms (see Tables S1 for all sites and habitats for each group of organisms). Then, we averaged all these separate samples to produce a single mean simulated sample. This was exactly the same procedure used to obtain the mean observed SAD at each scale as described in the main text. Thus, the simulated samples at site and habitat scales actually represents the mean of separate samples generated from each sites or habitats.

To perform the NB simulated samples described in the main text of this manuscript, the values of the overdispersion parameter κ for each species at each scale were estimated based on the empirical data using maximum likelihood techniques. Estimates of κ at site scale and for rare species generally failed to converge. In these cases, we assigned an arbitrary value of κ that was equal to the community average of the parameter at each scale. Therefore, NB draws were generated using species and scale-specific values of κ, an approach which closely mimics what is observed in nature. The only exception was the estimation of κ at site scale that had only three or four observations for the three groups of organisms. In this particular case, although the number of observations per species was low, we assumed that they followed a NB distribution. Thus, we randomly sampled abundance values from the empirical data to create a random NB regional SAD. This procedure guarantees that the scale of observation is preserved (the abundance of each species can come from any site) and that on average the size of the community is $n_{site}/N$ times smaller than $N$.

The generation of simulated samples of species biomass distribution was conducted using the same procedures described above but with an additional step. After generating the draw $v$, which is a vector of abundances, the value of abundance of each species was used to conduct a sample with replacement from the empirical values of biomass of the same species at the same scale. If the species abundance was zero, a biomass value equals zero was assigned to the species. This additional step was easy to implement but it certainly generated some unrealistic situation for rare species – since these species have few observed biomass values, sampling may have generated unlikely draws of populations with identical biomass values. However, we believe that such draws have not changed the overall outcomes of the simulation. We could have tried to fit continuous distributions to empirical biomass data and sample values from these fitted distributions but this would add more complexity to the analysis, related to the arbitrary selection of a statistical distribution and to the estimation of its parameters using few observations at some scales or for some species. Therefore, the species biomass distributions were simulated using the general codes:

```
# Defining the empirical biomass distributions of each species
      data1 <- tapply (data$biomass, list(data$sp), function(x) x)
# Sampling with replacement biomass values from the empirical distribution
      sim.data = NULL
      for (i in 1:length(data1)) {
            if (v[i]==0) { value= 0 } else {
                  value= sum(sample(x= data1[[i]], size= v[i], rep=TRUE)) }
            if (is.null(sim.data)) { sim.data= value } else {
                  sim.data= c(sim.data, value) }
      }
```

In the example above, the sampled individuals of each species are represented by the rows of a data frame arbitrarily named 'data' that are specified by columns 'biomass' (empirical biomass observations) and 'sp' (name of the species to which the individual belongs). The loop guarantees that the number of biomass values sampled for each species matches the abundance of the same species in the vector of abundance *v*. This procedure accepts expected abundances simulated using both Poisson and BN distributions.

Notice that we decided to use an approach similar to parametric bootstrapping *sensu* Manly (2006). Another way to test if the change in the shape of SAD is only a matter of sample size would be to use the analytical framework that takes into account the effect of sampling intensity to model SADs (Bulmer 1974; Green & Plotkin 2007). Based on sampling theory, this framework has explicit predictions of how the shape of sampled SAD should change according to the sampling intensity ($N_{\text{sample}}/N_{\text{metacommunity}}$), also known as sampling ratio (Dewdney 1998). However, we decided for our approach because there is no need to assume an underlying SAD model, such as a Lognormal or Gamma distributions, since all the information needed for generating the draws comes from the observed SADs themselves. In addition, there is no need to assume any size of an unknown metacommunity. Although the use compound models of SADs was recently facilitated by implementation of specific software solutions (*e.g.* Poisson Lognormal – Engen *et al.* 2002), there is no such ease for aggregate sampling and trying to fit a NB Lognormal using numerical integration usually leads to an impracticable amount of optimization errors (Foster & Warton 2007). Moreover, both approaches suffer from the same limitations when dealing with biomass (continuous data) instead of counts of individuals. It should be notice, however, that both approaches are quite similar in their basic principles, in particular the existence of a sampling intensity ‘control’ on the SAD shape at each scale. Consequently, the analysis using the Poisson Lognormal for the beetle and bird datasets ended up in the same qualitative results presented in the main text (results not shown).

**Appendix S3**. Comparison between observed and simulated SADs for trees, beetles and birds sampled at different scales, but using the same mean sample size.

**Table S2**. Summary of the Kolmogorov-Smirnov tests comparing the observed and simulated SADs for counts of individuals. Values correspond to statistic and p-value (separate by semicolons) of the test comparing the mean observed SAD and the mean simulated SADs for each reference sampling scale. Values in bold mean that observed and simulated SADs are different (α= 5%). For each reference sampling scale (*i.e.*, regional, site and habitat), we simulated communities using the same number of species and relative densities but with a mean sample size of the corresponding scale of simulation (*i.e.*, site, habitat and sample − see Online Resource 2). Simulations assumed conspecific aggregation (Negative binomial sampling) for trees and complete randomness (Poisson sampling) for birds and beetles.

| **Simulation scale** | **Organism** | **Reference scale** | | |
|---|---|---|---|---|
| | | Regional | Site | Habitat |
| Site | Trees | 0.088; 0.292 | | |
| | Beetles | 0.166; 0.624 | | |
| | Birds | 0.098; 0.759 | | |
| Habitat | Trees | 0.038; 0.999 | 0.063; 0.935 | |
| | Beetles | 0.151; 0.857 | 0.118; 0.976 | |
| | Birds | 0.123; 0.774 | 0.201; 0.147 | |
| Sample | Trees | 0.089; 0.994 | **0.264; 0.033** | **0.321; 0.004** |
| | Beetles | 0.190; 0.945 | 0.159; 0.990 | 0.167; 0.979 |
| | Birds | 0.124; 0.974 | 0.202; 0.511 | **0.333; 0.034** |

**Table S3**. Summary of the Kolmogorov-Smirnov tests comparing the observed and simulated SADs for biomass. Values correspond to statistic and p-value (separate by semicolons) of the test comparing the mean observed SAD and the mean simulated SADs for each reference sampling scale. See caption of Table S2 and Online Resource 2 for further details on the simulation procedures.

| **Simulation scale** | **Organism** | **Reference scale** | | |
|---|---|---|---|---|
| | | Regional | Site | Habitat |
| Site | Trees | 0.046; 0.969 | | |
| | Beetles | 0.102; 0.952 | | |
| | Birds | 0.055; 0.996 | | |
| Habitat | Trees | 0.135; 0.117 | 0.056; 0.955 | |
| | Beetles | 0.129; 0.902 | 0.152; 0.737 | |
| | Birds | 0.107; 0.792 | 0.124; 0.552 | |
| Sample | Trees | 0.21; 0.092 | 0.136; 0.275 | 0.142; 0.178 |
| | Beetles | 0.112; 0.986 | 0.126; 0.962 | 0.166; 0.727 |
| | Birds | 0.113; 0.933 | 0.155; 0.574 | 0.257; 0.051 |

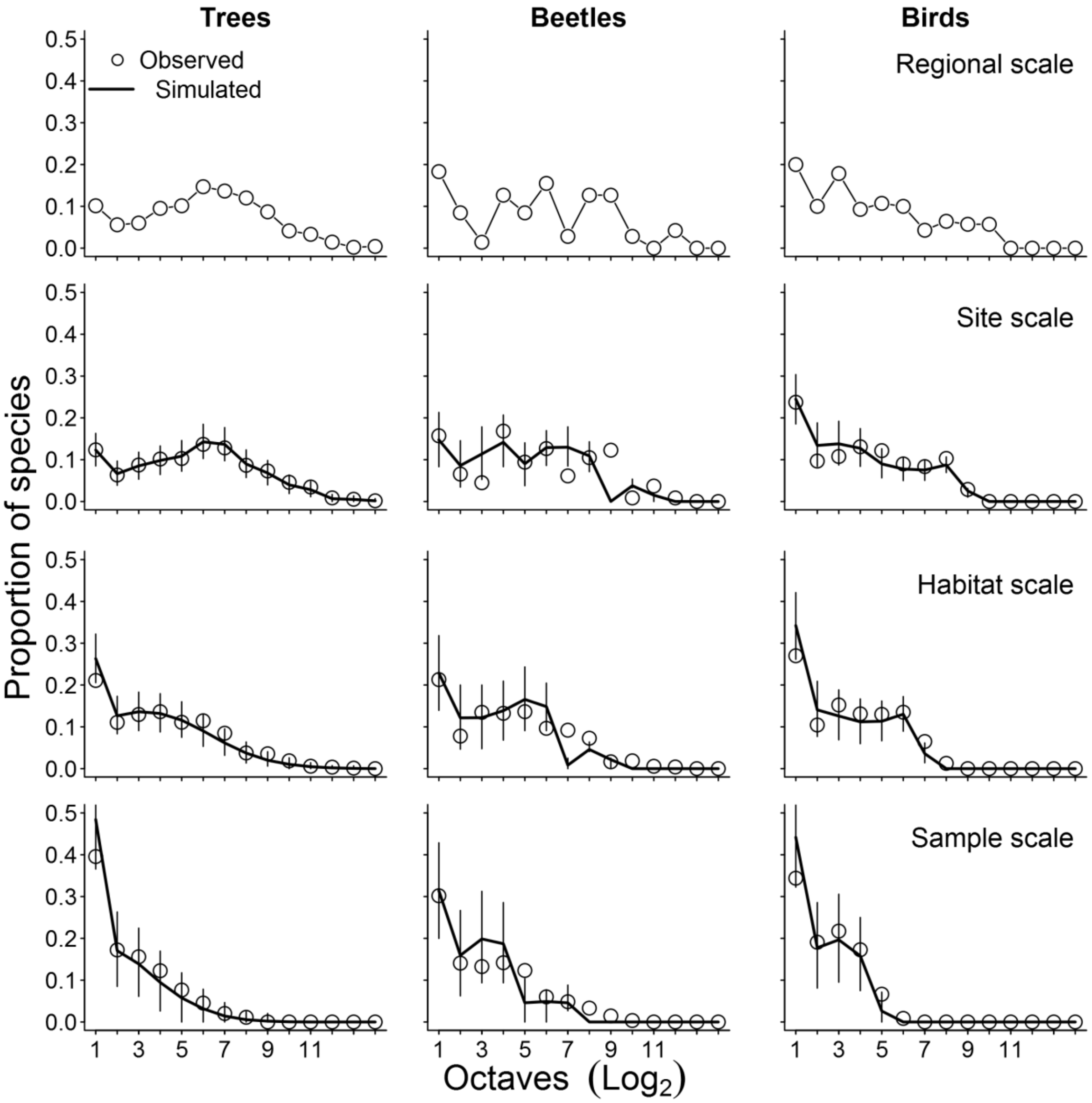


**Fig. S1**. Preston plots of the observed (circles) and simulated (thick lines) species abundance distributions (SAD) constructed using counts of individuals. Values presented are the mean proportion of species in each octave of abundance for trees, beetles and birds at four sampling scales. First octave contains species with less than $2^1$ individuals, second octave contains species with $\geq 2^1$ and $< 2^2$ individuals, and so on. These plots were produced using the exactly same data used to produce Figure 1 of the main text. Vertical thin lines depict the 90% quantiles of expected simulated values.

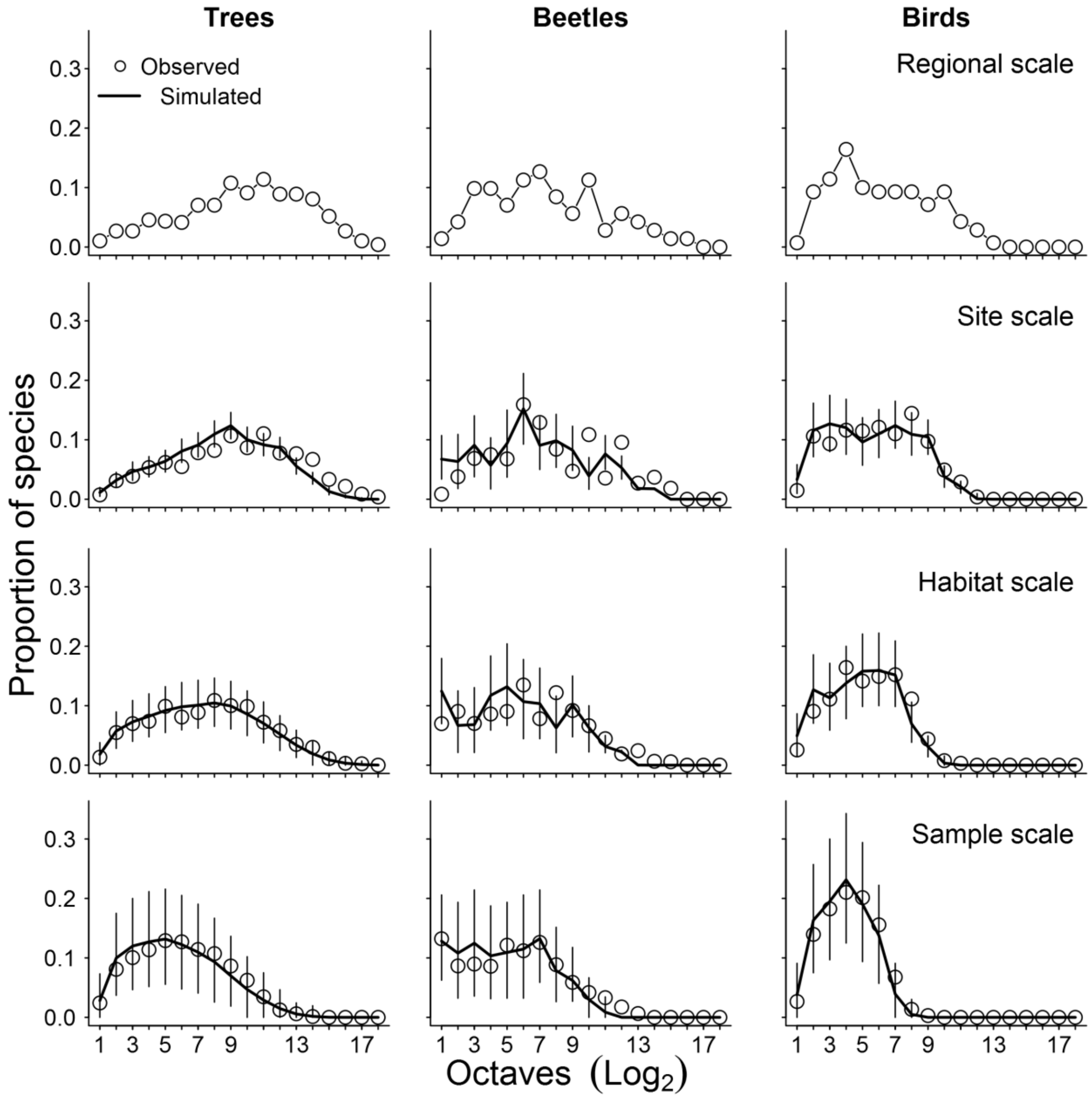


**Fig. S2**. Preston plots of the observed (circles) and simulated (thick lines) species abundance distributions (SAD) constructed using biomass. Values presented are the mean proportion of species in each octave of abundance for trees, beetles and birds at the four sampling scales. But here the first octave contains species with biomass less than $2^{-6}$ kg, second octave contains species with $\geq 2^{-6}$ and $< 2^{-5}$ kg, and so on. These plots were produced using the exactly same data used to produce Figure 2 of the main text. Vertical thin lines depict the 90% quantiles of expected simulated values.

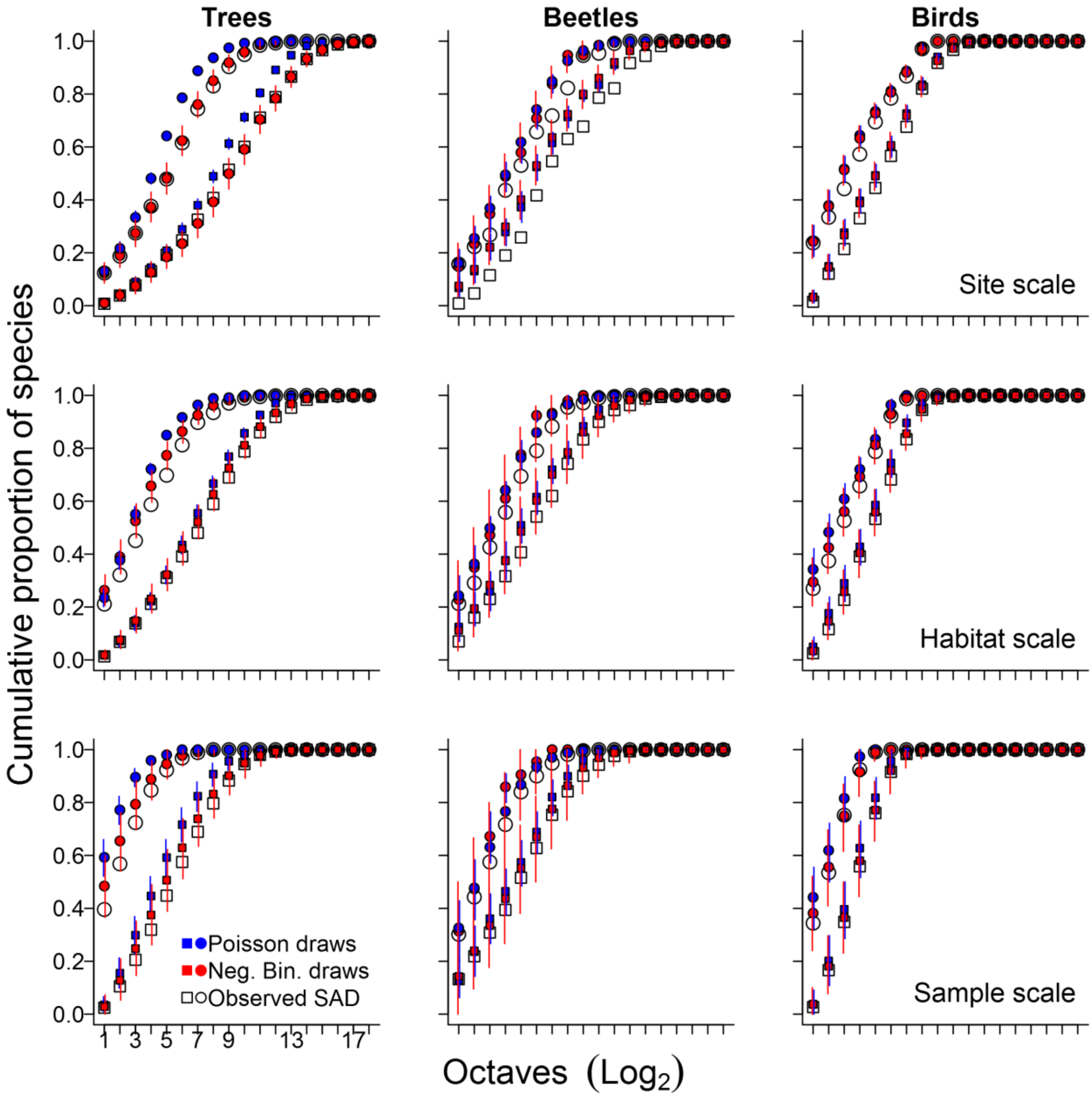


**Fig. S3**. Comparison between observed (white symbols) and simulated (colored symbols) mean cumulative proportions of species in each octave of abundance for trees, beetles and birds at different scales. Each panel contains the observed proportions for counts of individuals (circles) and biomass (squares) and simulated proportions obtained using Poisson (blue circles and squares) and Negative Binomial sampling (red circles and squares). The results in these graphics were obtained in the same way as those presented in Figures 1 and 2 in the main text, but here we use cumulative proportions per octave instead of mean proportions. The comparison between expected and observed proportions is made based on the 90% quantiles of the simulated values (vertical lines).

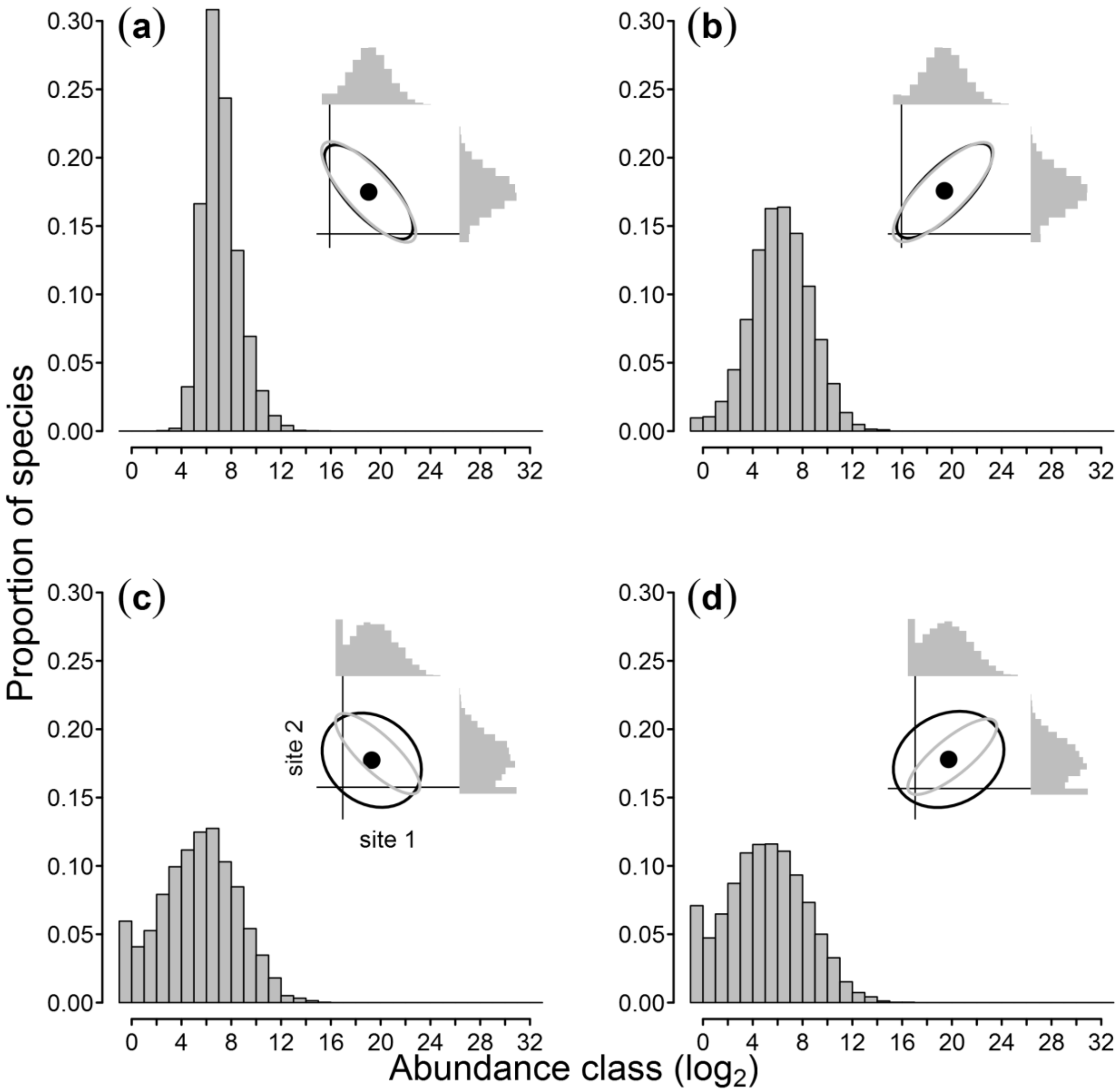


**Fig. S4**. The influence of conspecific aggregation and of the correlation of species abundances among communities (beta diversity) in the shape of the SAD, while combining samples to increase the sampling scale. Here, we exemplify this influence by pooling together two samples taken from a bivariate Lognormal distribution with the exact same parameters and controlling for possible sample size effects. We present the resulting Preston plots from the combination of sites where individuals of all species are randomly distributed and beta diversity is high (a) or low (b) and when there is conspecific aggregation with high (c) or low beta diversity (d). In each panel, the inset plot shows the ellipses that encompass 95% of the bivariate distributions and the marginal Preston plots of the sample SADs. Gray ellipses delimit the joint distribution of abundances in the communities and black ellipses delimit the distribution of abundances in the samples.

**Appendix S4**. Influence of conspecific aggregation and beta diversity in the shape of the SAD. Codes are given in language R (http://www.r-project.org).

Here, we illustrate how to simulate two samples from the same underlying Log-normal metacommunity under different scenarios of conspecific aggregation and beta diversity (i.e. the correlation of species abundances between communities). We use a parametric simulation approach by taken samples from a multivariate Log-normal. The main advantage of this approach relies on the existence of a parameter to define the exact correlation of species abundances between communities, used here as a measure of the beta diversity. As a disadvantage, the approach is restricted to a Log-normal underlying SAD.

The basic steps to perform this simulation are the following. We first define the number of metacommunity replicates with a pool of species of size 'S'. For each replicate, we draw the logarithm expected abundances for each species from a multivariate Normal distribution with parameters 'mu' and 'sd' and with the matrix 'rho' of correlation between observations (i.e. species abundances). This multivariate Normal distribution has dimension equal to the number of replicates chosen, which in this example will be two, for simplicity. Then, each metacommunity replicate is sampled using the Negative Binomial and Poisson distributions, to simulate a sampling process with and without conspecific aggregation, respectively.

In the accompanying R script (‘supplement_functions.R’), we provide the function *'rsad.mvln'* to perform these simulations. The function is documented in roxygen2 syntax (Whickham et al. 2017) and include as an example the codes to generate the data presented in Figures 3 and Fig S4. The script has also the functions *'p0'* and *'p1'* to generate these two figures, along with the scripts to reproduce the figures as examples in the function documentation.

**Appendix S5**. Alternative ways to obtain Preston plots at different sampling scales.

There are different ways to obtain Preston plots of pooled samples taken from different sampling scales. In Fig. S5a, the species abundance distribution (SAD) at a higher scale is depicted as a Preston plot of the abundances of species from pooled samples. The SAD at the scale below is depicted in three alternative ways: Averaged Preston Plot (Fig. S5b, the proportion of species in each sample that fall in each octave is averaged across samples); Preston plot of the sample mean abundances of each species (Fig. S5c); and as an Averaged Preston plot smoothed by Preston's binning method (Fig. S5d - Preston 1948). The apparent change in shape from panels a to b (or d) vanishes if we use mean abundances (panel c) instead of average number of species in Preston plots. By doing so, rare species just shift to lower octaves as we downscale. In contrast, averaged Preston plots implicitly sets a minimum octave to where rare species are packed with downscaling, changing the shape of the plot. Therefore, Preston plots of mean abundances partition out the effects of increasing sample size by pooling non-overlapping sampling units. A more direct way to do this is to simulate expected mean number of species by keeping sample size constant (Figures 1 and 2 on main text).

The number of instances sampled (sites or time occasions) obviously affects the accuracy of the estimated mean species abundances and mean proportion of species in each octave. To produce the data presented in Fig. S5, we used the function *'rsad.mvln'* described above (Online resource 4) and took random draws from a 100-variate Poisson-Lognormal distribution to simulate species abundances from 100 instances, with no spatial aggregation of conspecific individuals. Parameters used in the draws were: 10,000 Poisson draws from a 100-variate normal with parameters mean of log abundances $\mu_i$= 5, standard deviation of log abundances $\sigma_{ii}$=3.5 and species abundance correlation among communities $\rho_{ij}$=0.8. Hence, the sampled 100-variate normal has a 100×100 covariance matrix with diagonal elements $\sigma^2_{ii}$= 12.25 and off-diagonal elements $\sigma^2_{ij}$= 9.8. We provide the function *'ave.preston'* which take as output the community and sample matrices with the abundances of species at each site and returns the counts of species at each Preston octave using the two averaging methods discussed in this section. The example in the documentation of this function reproduces Figure S5.

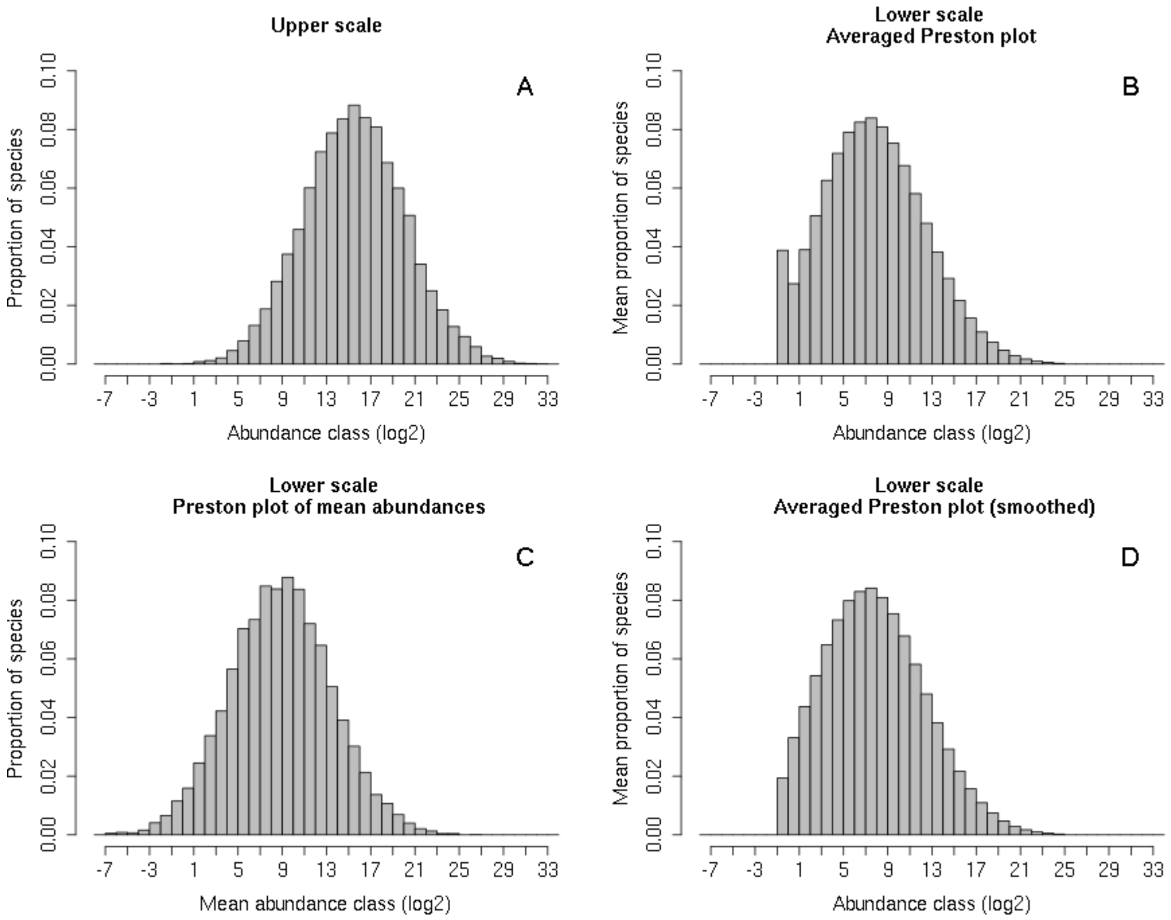


**Fig. S5**. The veil effect is a property of the averaged Preston plot. Panel A shows the abundance distribution at an upper scale, which is the sum of abundances of the abundances of species at 100 sites (or moments). Panels B-D show alternative ways to show the average abundance distributions at the lower (site) scale. In the averaged Preston Plot (B) the proportion of species in each abundance class (octaves) is the mean of proportions over sites. Panel C shows a Preston plot of the mean abundances of species over sites. Panel D shows the averaged Preston using Preston's binning method, which results in the well-known left-truncated or 'veiled' shape.
