## Supporting R Codes for "Disentangling the effects of sampling scale and size on the shape of species abundance distributions"

## Disentangling the effects of sampling scale and size on the shape
## of species abundance distributions.
## Renato Lima et al.
## R functions used in supplementary material.
## Functions documented in Roxygen syntax. Examples reproduce
## simulations and plots of the paper and supplements.

## Dependencies
require(mvtnorm)
require(sads)
require(car)

#' Poisson and Negative binomial samples of a multivariate lognormal
#'
#' This function simulates samples from communities at multiple
#' sites. The abundances of species in each site are
#' drawn from a multivariate lognormal. The samples take a fraction of
#' the individuals at each site, with any degree of spatial clumping
#' at the sampling unit.
#'
#' @param S positive integer. Total number of species in the metaccomunity.
#' @param replicates positive integer. Number of sites
#' @param mu vector of reals. Parameter of each marginal distribution
#'     of the multivariate lognormal. mu is the expected value of the
#'     logarithms of abundances at each site. If a single value is
#'     provided, all sites are set with the same parameter value.
#' @param sigma vector of reals. Parameter of each marginal distribution
#'     of the multivariate lognormal. sigma is the standard deviation of the
#'     logarithms of abundances at each site. If a single value is
#'     provided, all sites are set with the same parameter value.
#' @param rho matrix of reals. Pairwise matrix of Pearson correlations
#'     of abundances between pairs. Diagonals must be one, as they
#'     express the correlations of abundances of each site to
#'     itself. If a single value is provided, a 
#'     matrix with the same value for all off-diagonal elements.
#' @param frac real 0 < frac >=1. The sampling effort, which is the
#'     proportion of individuals in each site that are included in the
#'     sample.
#' @param pois.samp logical, should random sample of individuals be
#'     used? If TRUE a Poisson sample of 'frac' individuals is drawn
#'     ate each site. If FALSE a Negative Binomial sample is drawn
#'     instead, to simulate a sample of clumped inidividuals.
#' @param k positive real. The dispersion coefficient, which is the
#'     parameter 'size' of the negative binomial distribution (see
#'     'dnbinom'). The smaller the value of k the more clumped are
#'     individuals ate the scale of sampling units. 
#' @param \ldots further parameters to be passed to rmvnorm.
#' @return a list with two species by site matrices, both with 'S'
#'     rows and 'replicate columns'. Matrix 'commmunity' has the abundances
#'     of each species at each site in the metacommunity, as drawn
#'     from the multivariate lognormal defined by 'mu', 'sigma' and
#'     'rho'. Matrix 'sample' has the number of individuals of each
#'     species in the samples drawn at each site.
#' @examples
#' Simulated data used in Figures 3 and S4
#' set.seed(42)
#' ## 1. Bivariate lognormal with negative correlation
#' ## Poisson samples
#' sim1.p <- rsad.mvln(S = 1e4, replicates = 2, frac = 1,
#'                     mu = 3.5, sig = 1.7, rho = -0.8)
#' ## Negative binomial samples
#' sim1.nb <- rsad.mvln(S = 1e4, replicates = 2, frac = 1, 
#'                      mu = 3.5, sig = 1.7, rho = -0.8,
#'                      pois.samp = FALSE, k = 0.2)
#' ## 2. Bivariate lognormal with positive correlation
#' ## Poisson samples
#' sim2.p <- rsad.mvln(S = 1e4, replicates = 2, frac = 1,
#'                     mu = 3.5, sig = 1.7, rho = 0.8)
#' ## Negative binomial samples
#' sim2.nb <- rsad.mvln(S = 1e4, replicates = 2, frac = 1,
#'                      mu = 3.5, sig = 1.7, rho = 0.8,
#'                      pois.samp=FALSE, k = 0.2, )
rsad.mvln <- function(S, replicates, mu, sig, rho, frac = 1, pois.samp=TRUE, k, ...){
    if(length(mu)==1) mu <- rep(mu, replicates)
    if(length(sig)==1) sig <- rep(sig, replicates)
    if(length(rho)==1){
        rho <- matrix(rho, ncol=replicates, nrow=replicates)
        diag(rho) <- 1
    }
    sig.mat <- outer(sig,sig,"*")
    cov.mat <- sig.mat*rho
    com <- samp <- matrix(nrow=S, ncol=replicates)
    com <- rmvnorm(n=S, mean=mu, sigma=cov.mat, ...)
    if(pois.samp){
        for(i in 1:replicates)
            samp[,i] <- rpois(n=S, lambda=exp(com[,i])*frac)
    }
    else{
        if(length(k)==1) k <- matrix(k, nrow=S, ncol=replicates)
        else if(length(k)==S) k <- matrix(k, ncol=replicates)
        for(i in 1:replicates)
            samp[,i] <- rnbinom(n=S, size=k[,i], mu=exp(com[,i])*frac)
    }
    list(community=exp(com), sample=samp)
}

#' Averaged and mean abundance Preston tables (Fig. S5)
#'
#' Given a species x sites matrix of abundance of species in the whole
#' metacommunity and in samples taken at each site, calculates the
#' mean proportion of species in each abundance class over all sites
#' and also the mean abundance of each species per site.
#' @param samp a species x sites matrix with the number of individuals
#' of each species sampled at each site.
#' @param com a species x sites matrix with the abundance of
#' individuals of each species at each site.
#' @param oc vector of octave (abundances classes in log2 scale) to
#' use to tally the species. See 'sads::octav'
#' @param preston logical; should Preston's binning method be used to
#' tally species in octaves? See 'sads::octav'.
#' @return a list with three objects of class 'octav'. Each object is
#' a data.frame with the number of species in each abundance
#' octave. See 'sads::octav'. Object 'com' is the octave table for the
#' species abundances in the metacommunity. Object 'sample.ab.mean' is
#' the octave table for the mean abundance of species per site. Object
#' 'sample.sp.mean' is the table of mean number of species in each
#' abundance class over sites.
#' @examples
#' ## Figure S5
#' ## Simulate a metacommunity with 1e4 species and 100 sites
#' ## and a sample of 0.1 of the individuals at each site
#' set.seed(42)
#' sim3 <- rsad.mvln(S = 1e4, replicates = 100, frac=1,
#'                   mu = 5, sig=3.5, rho = 0.8)
#' ## Preston tables
#' sim3.oc <- ave.peston(sim3$sample, sim3$community)
#' ## With Preston's binning method
#' sim3b.oc <- f5(sim3$sample, sim3$community, preston=TRUE)
#' ## Plot of figure S5
#' par(mfrow=c(2,2), cex.axis=1.25, cex.lab=1.3, cex.main=1.25)
#' plot(sim3.oc$com, prop=TRUE, x.oct=TRUE, ylim=c(0,0.1), main="Upper scale")
#' text(31,.095, expression(bold((a))), cex=2)
#' plot(sim3.oc$sample.sp.mean, prop=TRUE, x.oct=TRUE, ylab="Mean proportion of species", ylim=c(0,0.1),
#'      main="Lower scale \n Averaged Preston plot ")
#' text(31,.095, expression(bold((b))), cex=2)
#' plot(sim3.oc$sample.ab.mean, prop=TRUE, x.oct=TRUE, xlab="Mean abundance class (log2)", ylim=c(0,0.1),
#'      main="Lower scale \n Preston plot of mean abundances")
#' text(31,.095, expression(bold((c))), cex=2)
#' plot(sim3b.oc$sample.sp.mean, prop=TRUE, x.oct=TRUE, ylab="Mean proportion of species", ylim=c(0,0.1),
#'      main="Lower scale \n Averaged Preston plot (smoothed)")
#' text(31,.095, expression(bold((d))), cex=2)
ave.preston <- function(samp, com, oc, preston=FALSE){
    s.ab.mean <- apply(samp,1,mean)
    c.ab.sum <- apply(com,1,sum)
    if(missing(oc)){
        oc.range <- apply(cbind(samp,s.ab.mean, c.ab.sum),2,af1)
        oc <- min(oc.range):max(oc.range)
    }
    spcount <- apply(samp, 2, af2, oc, preston=preston)
    s.ab.mean.oc <- octav(s.ab.mean, oc)
    c.ab.sum.oc <- octav(c.ab.sum, oc)
    df1 <- data.frame(oc, 2^oc, apply(spcount,1,mean))
    names(df1) <- names(s.ab.mean.oc)
    sp.mean <- new("octav", df1) 
    list(com=c.ab.sum.oc, sample.ab.mean=s.ab.mean.oc, sample.sp.mean=sp.mean)
}

## Plotting functions 

#' Averaged rank-abundance plot with bivariate Poisson-lognormal
#' distribution in inset
#'
#' This function generates the plots shown in each panel of Figure 3
#' @param X vector of number of individuals of each species sampled in
#'     site 1
#' @param Y vector of number of individuals of each species sampled in site 1
#' @param community species by site matrix of abundances of each
#'     species in each site.
#' @param max.oc maximum number of octaves to be used in Preston's plots.
#' @param fig the NDC coordinates for the plot, see help(par).
#' @param xlab2 x label for the inset plot.
#' @param ylab2 y label for the inset plot.
#' @param label plot label to be included at top right croner with 'mtext'
#' @param \ldots further parameters to be passed to 'plot'.
#' @examples
#' ## Figure 3 of Lima et al.
#' def.par <- par(no.readonly = TRUE)
#' par(def.par, fig=c(0,1,0,1)); plot(1,1, type="n", axes=FALSE,xlab="",ylab="")
#' p0(X=sim1.p$sample[,1],Y=sim1.p$sample[,2],
#'      community=sim1.p$community,  fig=c(0,1/2,1/2,1), xlab2="",
#'      ylab2="", label=expression(bold((a))), ylim=c(2.5e-07,0.025),
#'      xlim=c(0.75,10000),xlab="", ylab="") 
#' p0(X=sim1.nb$sample[,1],Y=sim1.nb$sample[,2],
#'      community=sim1.nb$community,  fig=c(1/2,1,1/2,1), xlab2="",
#'      ylab2="", label=expression(bold((b))), ylim=c(2.5e-07,0.025),
#'      xlim=c(0.75,10000),xlab="", ylab="")
#' p0(X=sim2.p$sample[,1],Y=sim2.p$sample[,2],
#'    community=sim2.p$community, fig=c(0,1/2,0,1/2), xlab2="", ylab2="",
#'    label=expression(bold((a))), ylim=c(2.5e-07,0.025),
#'    xlim=c(0.75,10000),xlab="", ylab="")
#' p0(X=sim2.nb$sample[,1],Y=sim2.nb$sample[,2],
#'    community=sim2.nb$community, fig=c(1/2,1,0,1/2), xlab2="",
#'    ylab2="", label=expression(bold((b))), ylim=c(2.5e-07,0.025),
#'    xlim=c(0.75,10000),xlab="", ylab="")
#' par(fig=c(0,1,0,1), new=T)
#' mtext("Species relative abundance", side=2, line=-1.25, cex=1.5)
#' mtext("Abundance rank", side=1, line=-1.5, cex=1.5)
p0 <- function(X, Y, community, max.oc=27, fig, xlab2="", ylab2="", label="", ...){
    x <- log2(X)
    y <- log2(Y)
    com <- log2(community)
    ## Confidence ellipse
    e1a <- dataEllipse(x[X>0&Y>0],y[X>0&Y>0], draw=FALSE, levels=0.95)
    ## Breaks and xlim base on ellipse limits
    b.x <- floor(min(c(e1a[,1],x[X>0]))):ceiling(max(c(e1a[,1],x[x>0])))
    b.y <- floor(min(c(e1a[,2],y[Y>0]))):ceiling(max(c(e1a[,2],y[Y>0])))
    ## tweaks to plot histograms nicely
    xhist <- hist(x, breaks=b.x, plot=FALSE)
    yhist <- hist(y, breaks=b.y, plot=FALSE)
    top <- max(c(xhist$counts, yhist$counts))
    xl <- abs(fig[1]-fig[2])
    yl <- abs(fig[3]-fig[4])
    ## Large plot: rank-abundance plot of averaged abundances
    par(las=1, fig=fig, new=TRUE, mgp = c(3.5,1,0), mar = c(5,5,4,4), cex.axis = 1.25, cex.lab=1.3)
    z <- X+Y
    par(mgp=c(2.1,0.3,0),mai=c(0.6,0.7,0.3,0.1),tcl=-0.2)
    plot(y=rad(z)$abund/sum(rad(z)$abund),x=rad(z)$rank, 
			xaxt="n",yaxt="n",log="xy",bty="l",pch=19,cex=0.75,cex.axis=0.75,...)
    axis(1,at=c(1,10,100,1000),lab=T,cex.axis=0.8)
    axis(2,at=c(0.0000001,0.000001,0.00001,0.0001,0.001,0.01),
         lab=c(0.0000001,0.000001,0.00001,0.0001,0.001,0.01),cex.axis=0.8)
    legend("topright",label, adj=c(0,0), cex=1.5, bty="n")	
    ## Inset plot
    ## Relative coordinates of insets
    fig2 <- c(fig[1]+xl*.225,fig[1]+xl*.525,fig[3]+yl*.2,fig[3]+yl*.5)
    par(fig=fig2, mar = c(0,0,0,0), new=TRUE, las=0)
    plot(x,y,
         xlim=range(b.x),
         ylim=range(b.y),
         type="n",
         axes=FALSE
         )
    abline(v=0, h=0)
    mtext(xlab2, side=3, line=1.9)
    mtext(ylab2, side=4, line=1.8)
    ## Add 95% ellipses
    dataEllipse(x[X>0&Y>0],y[X>0&Y>0], levels=0.95, col="black", plot.points=FALSE, add=TRUE, center.cex=1)
    dataEllipse(com, levels=0.95, col="gray", plot.points=FALSE, add=TRUE, center.pch=0, center.cex=1)
    ## The first histogram
    par(fig=c(fig2[2], fig2[2]+0.05, fig2[3], fig2[4] ), new=TRUE, mar = c(0.1,0,0,0))
    barplot(yhist$counts, axes = FALSE, xlim = c(0, top),
            space = 0, horiz = TRUE, border=NA)
    ## The second histogram
    par(fig=c(fig2[1], fig2[2], fig2[4], fig2[4]+0.05), new=TRUE, mar = c(0.1,0,0,0))
    barplot(xhist$counts, axes = FALSE, ylim = c(0, top),
            space = 0, horiz=FALSE, border=NA)
}


#' Averaged Preston plot with bivariate Poisson-lognormal
#' distribution in inset
#'
#' This function generates the plots shown in each panel of Figure S4
#' @param X vector of number of individuals of each species sampled in
#'     site 1
#' @param Y vector of number of individuals of each species sampled in site 1
#' @param community species by site matrix of abundances of each
#'     species in each site.
#' @param max.oc maximum number of octaves to be used in Preston's plots.
#' @param fig the NDC coordinates for the plot, see help(par).
#' @param xlab2 x label for the inset plot.
#' @param ylab2 y label for the inset plot.
#' @param label plot label to be included at top right croner with 'mtext'
#' @param \ldots further parameters to be passed to 'plot'.
#' @examples
#' ## Figure S4 of Lima et al.
#' def.par <- par(no.readonly = TRUE)
#' par(def.par, fig=c(0,1,0,1)); plot(1,1, type="n", axes=FALSE,xlab="",ylab="")
#' p1(X=sim1.p$sample[,1],Y=sim1.p$sample[,2],
#'      community=sim1.p$community,  fig=c(0,1/2,1/2,1), xlab2="",
#'      ylab2="", label=expression(bold((a))), ylim=c(0,.3), xlab="", ylab="") 
#' p1(X=sim1.nb$sample[,1],Y=sim1.nb$sample[,2],
#'      community=sim1.nb$community,  fig=c(1/2,1,1/2,1), xlab2="",
#'      ylab2="", label=expression(bold((b))), ylim=c(0,0.3), xlab="", ylab="")
#' p1(X=sim2.p$sample[,1],Y=sim2.p$sample[,2],
#'    community=sim2.p$community, fig=c(0,1/2,0,1/2), xlab2="", ylab2="",
#'    label=expression(bold((a))), ylim=c(0,0.3), xlab="", ylab="")
#' p1(X=sim2.nb$sample[,1],Y=sim2.nb$sample[,2],
#'    community=sim2.nb$community, fig=c(1/2,1,0,1/2), xlab2="",
#'    ylab2="", label=expression(bold((b))), ylim=c(0,0.3), xlab="", ylab="")
#' par(fig=c(0,1,0,1), new=T)
#' mtext("Species relative abundance", side=2, line=-1.25, cex=1.5)
#' mtext("Abundance rank", side=1, line=-1.5, cex=1.5)
p1 <- function(X, Y, community, max.oc=27, fig, xlab2="", ylab2="", label="", ...){
    x <- log2(X) 
    y <- log2(Y)
    com <- log2(community)
    ## Confidence ellipse
    e1a <- dataEllipse(x[X>0&Y>0],y[X>0&Y>0], draw=FALSE, levels=0.95)
    ## Breaks and xlim base on ellipse limits
    b.x <- floor(min(c(e1a[,1],x[X>0]))):ceiling(max(c(e1a[,1],x[x>0])))
    b.y <- floor(min(c(e1a[,2],y[Y>0]))):ceiling(max(c(e1a[,2],y[Y>0])))
    ## tweaks to plot histograms nicely
    xhist <- hist(x, breaks=b.x, plot=FALSE)
    yhist <- hist(y, breaks=b.y, plot=FALSE)
    top <- max(c(xhist$counts, yhist$counts))
    xl <- abs(fig[1]-fig[2])
    yl <- abs(fig[3]-fig[4])
    ## Large plot: Preston plot of pooled abundances
    par(las=1, fig=fig, new=TRUE, mgp = c(3.5,1,0), mar = c(5,5,4,4), cex.axis = 1, cex.lab=1.3)
    z <- X+Y
    par(mgp=c(2.1,0.3,0),mai=c(0.6,0.7,0.3,0.1),tcl=-0.2)
    plot(octav(z, oct=0:max.oc), x.oct=TRUE, prop=TRUE, par.ax=list(lwd=1.5), ...)
    legend("topleft",label, adj=c(1,0), cex=1.5, bty="n")    
    ## Inset plot
    ## Relative coordinates of insets
    fig2 <- c(fig[1]+xl*.55,fig[1]+xl*.8,fig[3]+yl*.5,fig[3]+yl*.75)
    par(fig=fig2, mar = c(0,0,0,0), new=TRUE, las=0)
    plot(x,y,
         xlim=range(b.x),
         ylim=range(b.y),
         type="n",
         axes=FALSE
         )
    abline(v=0, h=0)
    mtext(xlab2, side=1, line=.2)
    mtext(ylab2, side=2, line=.3)
    ## Add 95% ellipses
    dataEllipse(x[X>0&Y>0],y[X>0&Y>0], levels=0.95, col="black", plot.points=FALSE, add=TRUE)
    dataEllipse(com, levels=0.95, col="gray", plot.points=FALSE, add=TRUE, center.pch=0)
    ## The first histogram
    par(fig=c(fig2[2], fig2[2]+0.05, fig2[3], fig2[4] ), new=TRUE, mar = c(0,0,0,0))
    barplot(yhist$counts, axes = FALSE, xlim = c(0, top),
            space = 0, horiz = TRUE, border=NA)
     ## The second histogram
    par(fig=c(fig2[1], fig2[2], fig2[4], fig2[4]+0.05 ), new=TRUE, mar = c(0,0,0,0))
    barplot(xhist$counts, axes = FALSE, ylim = c(0, top),
            space = 0, horiz=FALSE, border=NA)
}

## Accesory functions ##
af1 <- function(x){
    range(as.numeric(octav(x)$octave))
}

af2 <- function(x, oc, ...){
    octav(x, oct=oc, ...)$Freq
}
